# Cortical high-frequency oscillations (≈ 110 Hz) in cats are state-dependent and enhanced by a subanesthetic dose of ketamine

**DOI:** 10.1101/2023.05.31.543142

**Authors:** Santiago Castro-Zaballa, Joaquín González, Matías Cavelli, Diego Mateos, Claudia Pascovich, Adriano Tort, Mark Jeremy Hunt, Pablo Torterolo

**Author notes:** Corresponding author: Dr. Santiago Castro-Zaballa. Full postal address: General Flores Avenue 2125 P.C. 11800 Montevideo – Uruguay.

## Abstract

Ketamine is an NMDA receptor antagonist that has both antidepressant and anesthetic properties. At subanesthetic doses, ketamine can cause transient psychosis in humans, and is used to model psychosis in experimental animals. In rodents, subanesthetic doses of ketamine increase the power of high-frequency oscillations (HFO, 100-180 Hz) in the electroencephalogram and field potentials, a frequency band linked to cognitive functions. However, the effects of ketamine in higher mammals, with more translatable relevance, are poorly investigated. Here, we have examined cortical HFO during wakefulness, sleep, and after administering a sub-anesthetic dose of ketamine (15 mg/kg), utilizing the cat as an animal model. Four cats were implanted with cortical electrodes for chronic polysomnographic recordings. HFO power, connectivity, information flow directionality, and their relationships with respiratory activity were analyzed. During wakefulness, but not during sleep, we found that HFO were coupled with the inspiratory phase of the respiration. After ketamine administration, HFO were enhanced significantly and remained associated with the inspiratory phase. The analysis of the information flow after ketamine suggest that HFO originate from the olfactory bulb and stream towards the prefrontal cortex. Accordingly, occluding the nostrils significantly reduced HFO power in both the olfactory bulb and prefrontal cortex. In contrast, auditory stimulation did not affect HFO. In conclusion, spontaneous cortical HFO show certain state-dependent features in cats, and enhancement of this rhythm by ketamine may disrupt cortical information processing, which could contribute to some of the neuropsychiatric manifestations associated with ketamine.

**Highlights:**

- Ketamine is used to model psychosis in humans and experimental animals
- Subanesthetic doses of ketamine increase the power of high-frequency oscillations
- High-frequency oscillations are coupled with the inspiratory phase of respiration
- These oscillations originate in the olfactory bulb and stream to the neocortex
- Nostril occlusion lowers high-frequency activity in the olfactory bulb and neocortex

**Graphical abstract:** 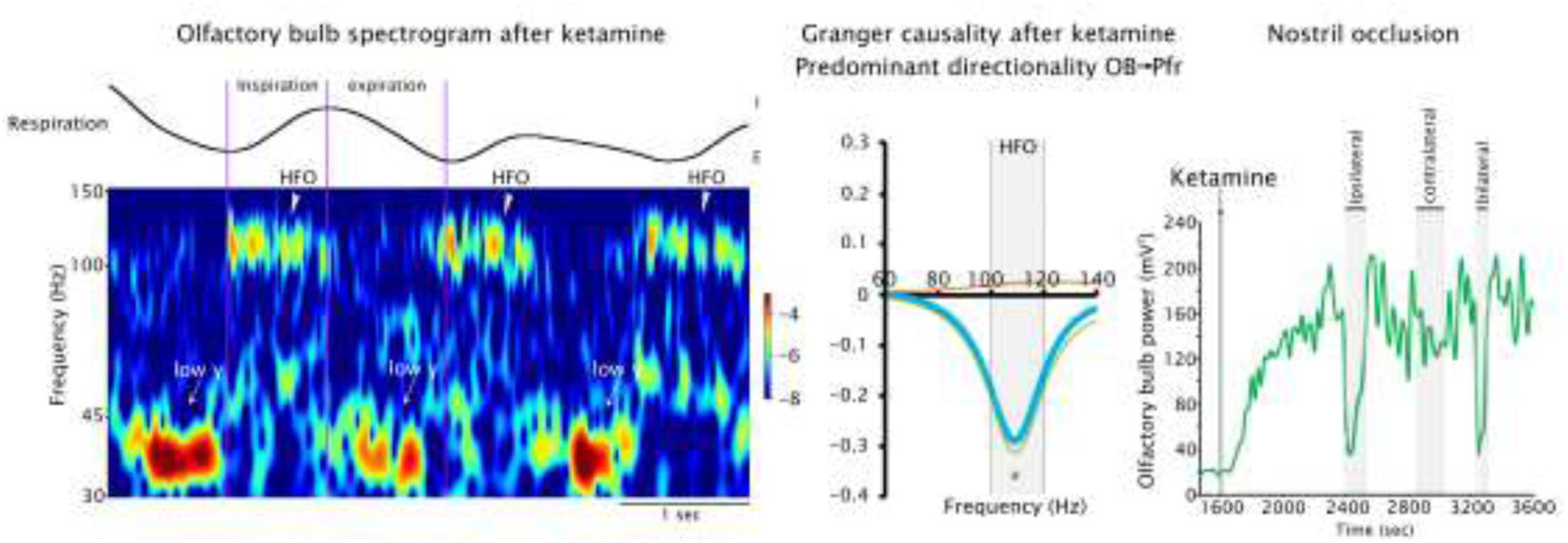

## 1. Introduction

High frequency oscillations (HFO > 100 Hz) recorded in the electroencephalogram (EEG), also known as fast gamma, are associated with cognitive functions such as decision-making and memory (Tort et al., 2008). In rodents, HFO are larger during wakefulness and REM sleep than non-REM (NREM) sleep (Hunt et al., 2009; Scheffer-Teixeira et al., 2013; Scheffzük et al., 2011), and their amplitude is modulated by theta oscillations in both the hippocampus and parietal cortex (Cavelli et al., 2018; González et al., 2020; Sirota et al., 2008; Tort et al., 2013). Furthermore, HFO coherence is greater during wakefulness than NREM sleep (Cavelli et al., 2018). In REM sleep, the synchronization patterns of HFO are complex, it depends on the phase of the theta cycle, and differ between inter- and intra-hemispheric derivates (Cavelli et al., 2018; González et al., 2020).

Ketamine is an N-methyl-D-aspartate (NMDA) receptor antagonist, that is used clinically in depression, pain and anesthesia (Diazgranados et al., 2010; Domino and Warner, 2010; Kohtala, 2021; Zarate et al., 2012). However, acute administration of ketamine at subanesthetic doses can lead to toxic psychosis, producing symptoms similar to those of schizophrenia (Beck et al., 2020; Krystal et al., 2005; Lahti et al., 1995). As such, subanesthetic doses of ketamine are used widely to model psychosis in humans and experimental animals (Frohlich and Van Horn, 2014).

Ketamine is known to produce complex effects on electrical oscillatory activity recorded in the EEG of both humans and rodents; for example, increases gamma power (30-100 Hz) (Akeju et al., 2016; Garwood et al., 2021; Pal et al., 2015). A growing body of evidence, mainly from rodents, has found marked increases in HFO power following subanesthetic doses of ketamine in a variety of cortical and subcortical areas, and the olfactory networks seem to be important in their generation (Cordon et al., 2015; Flores et al., 2015; Hunt et al., 2006, 2019; Hunt and Kasicki, 2013; Jones and Barth, 1999; Kealy et al., 2017; Nicolás et al., 2011a; Phillips et al., 2012; Ye et al., 2018). Further, this rhythm is sensitive to antipsychotics, indicating it may be a useful biomarker of ketamine psychosis (Olszewski et al., 2013). Importantly, recent evidence from scalp EEG recordings, has shown that subanesthetic ketamine may also increase HFO power in humans (Nottage et al., 2023). As such, it is necessary to understand better the potential networks involved in generating this rhythm using more translatable animal models.

Cats (*Felis silvestris catus*) share a similar gyrencephalic anatomy to humans and are thus suitable to probe the nature of cortical oscillatory activity in a translatable manner. Here, we examined HFO rhythms in the olfactory bulb (OB) and neocortical areas in cats chronically implanted with cortical EEG electrodes. The purpose of this study was two-fold: firstly, to examine changes in HFO over the sleep-wake cycle; and secondly, to examine the impact of a subanesthetic dose of ketamine on this rhythm.

## 2. Material and methods

### 2.1. Experimental animals

Four adult cats (2 neutered males of 6.5 kg, and 2 females of 3.5 and 5.4 kg) were obtained from and determined to be healthy by the institutional animal care facility. All experimental procedures were conducted in accordance with the Guide for the Care and Use of Laboratory Animals (National Research Council Committee, 2011), and were approved by the institutional animal care commission (Exp. N° 070153-000089-17). Adequate measures were taken to minimize animal pain, discomfort, or stress, and to use the minimum number of animals necessary to produce reliable scientific data.

### 2.2. Surgical procedures

The animals were chronically implanted with electrodes to monitor sleep and wakefulness. The surgical and experimental procedures were similar to those reported in our previous studies (Castro et al., 2013; Torterolo et al., 2016; Castro-Zaballa et al., 2019b; Cavelli et al., 2019; Pascovich et al., 2022). Before undergoing anesthesia, each cat was treated with atropine (0.04 mg/kg, intramuscularly, i.m.) and antibiotics (Tribrissen®, 30 mg/kg, i.m.). Anesthesia was induced using a combination of ketamine (15 mg/kg, i.m.) and xylazine (2.2 mg/kg, intramuscularly), and was maintained throughout the procedure with a mixture of isoflurane and oxygen (1-3%). The head was fixed in a stereotaxic frame, exposing the skull. Then, stainless-steel screw electrodes with a diameter of 1.4 mm were placed above the dura on the surface of various cortical regions, according to the atlas of (Berman and Jones, 1982). For this work, we analyzed recordings of the right hemisphere from dorsolateral prefrontal (Pfdl) and posterior parietal (Pp) cortices (4 animals). In 3 of these animals, recordings from the rostral prefrontal (Pfr) and OB were also performed and analyzed (Figure 1A).

**Figure 1.**
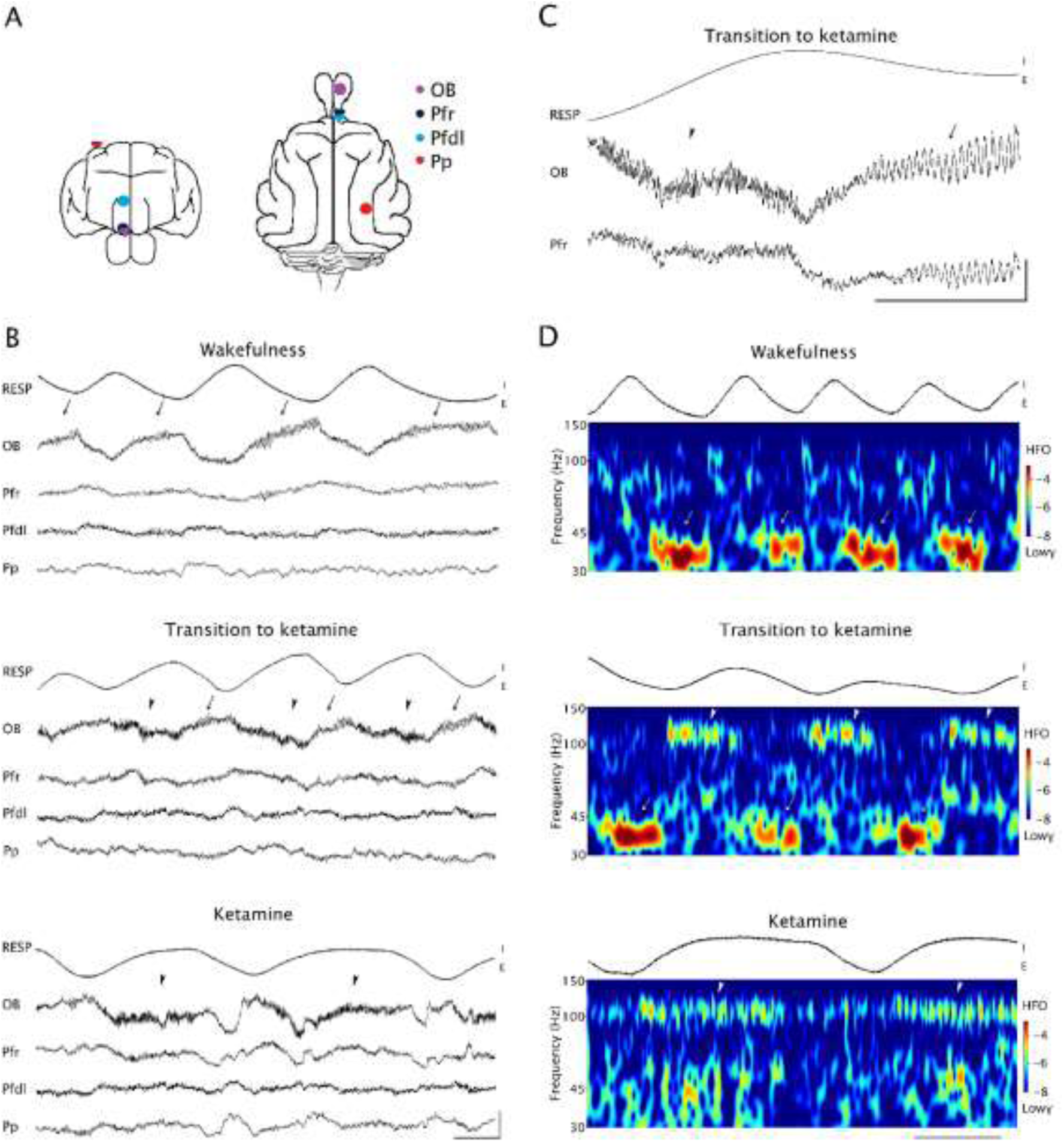
Spontaneous and ketamine-enhanced HFO and low-gamma oscillations in the cat EEG. A. Position of the recording electrodes on the surface of the cerebral cortex. The recordings were derived to a referential inactive electrode located on the left frontal sinus. B. Example of simultaneous raw recordings from a representative animal showing respiratory activity (Resp), and EEG from the olfactory bulb (OB), rostral prefrontal (Pfr), dorsolateral prefrontal (Pfdl) and parietal posterior (Pp) cortices during wakefulness, transition to ketamine and under the complete effect of ketamine. C. Simultaneous recordings during the transition to ketamine at higher magnification scale. D. Wavelet spectrograms from 30 to 150 Hz of the OB EEG of a representative animal during wakefulness, transition to ketamine and after ketamine; the respiratory activity is also shown. Arrows indicate low-gamma and arrowheads HFO “bursts” of activity. E, expiration, I, inspiration. Calibration bars: B, 1 mV and 0.5 sec; C, 0.1 mV and 0.5 sec; D, 1 sec.

The electrodes were connected to a Winchester plug and secured in place along with two plastic tubes, using acrylic cement. After the animals fully recovered from the surgical procedure, they were acclimated to the recording environment for a minimum of two weeks.

### 2.3. Experimental procedures

Experiments were conducted for 4 hours, between 11 A.M. and 3 P.M., in a temperature-controlled environment (21-23°C). During the acclimation period and the experiments, the head of the cat was secured in the stereotaxic position using four steel bars fitted into the chronically implanted plastic tubes. This allowed for fixing the head without causing discomfort while the body was kept in a sleeping bag.

The EEG (strictly electrocorticogram or intracranial EEG) was recorded using a monopolar (referential) configuration, with a common reference electrode placed in the left frontal sinus. Additionally, the electromyogram (EMG) of the nuchal muscles was recorded using a bipolar electrode placed on the skin over the muscle, the electrocardiogram was recorded using electrodes placed on the skin over the precordial region, and respiratory activity was recorded using a piezoelectric sensor located on the surface of the thorax. The sensor was calibrated both visually and through video, allowing for correlation with inspiration and expiration movements. The bioelectric signals were amplified × 1000 and filtered within the range of 0.1-500 Hz. The signals were then sampled at 1024 Hz with a 16-bit resolution and stored on a PC using Spike 2 software (Cambridge Electronic Design).

Data were collected during periods of spontaneous wakefulness, NREM sleep, and REM sleep, which were classified according to standard criteria (Ursin et al., 1981). In selected recording sessions, ketamine (15 mg/kg, i.m.) was administered at the beginning of the recordings. The cats were recorded daily for approximately 30 days in order to complete basal and treatment data sets.

Auditory stimulation (AS) was assessed in all the animals during both wakefulness and ketamine conditions. The AS lasted for 300 seconds and comprised of clicks with a duration of 0.1 ms, presented at ≍ 60 dB SPL, with a randomly varied frequency of presentation ranging from 1 to 500 Hz to prevent habituation (Castro et al., 2013; Castro-Zaballa et al., 2019b, 2019a). The sound stimuli were administered approximately 30 minutes into the baseline recording session (while the animals were in wakefulness) or 10 minutes after ketamine injection.

We ran a new set of experiments with nasal occlusion in the three animals with OB and Pfr electrodes. The animals were trained for several days before the *bona fide* experiment to tolerate the occlusion process, achieved by using cotton swabs moistened with saline solution to temporarily block their nostrils (Średniawa et al., 2021). However, only two of the three animals were able to fully adapt to the procedure. We carried out right nostril occlusion for 120 seconds, followed by a left nostril occlusion for the same duration, during both wakefulness and following ketamine administration (4 sessions per animal). After a 3 to 5-minute interval, we then performed bilateral nostril occlusion for 60 seconds.

### 2.4. Data analyses

#### 2.4.1. Power and connectivity

The analyses were similar to our previous reports (*op. cit*.). We obtained data from 16 baseline recording sessions (4 from each animal), where the animals slept *ad-libitum*, and 16 recording sessions under ketamine (4 from each animal). Sleep and wakefulness were quantified in 10-second epochs. In each recording, the EEG analysis was conducted on 12 windows of 100 seconds each, taken from artifact-free data of wakefulness, sleep, or ketamine (totaling 1200 seconds for each condition). Hence, the data bank consisted of 192 windows of EEG analysis (12 windows × 16 recordings) for wakefulness, NREM, REM sleep and ketamine in Pfdl and Pp, and 144 windows (12 × 12) for the OB and Pfr analysis.

Based on the results of our preliminary studies during ketamine, which showed a peak in HFO power between 100 and 120 Hz, this study focused on the analysis of this specific frequency band. For each selected window, the magnitude squared coherence was analyzed between two channels using the COHER 1S script in Spike 2 software. The 100-second period of analysis was divided into 100 blocks with a sample rate of 1024 Hz and a bin size of 2048 samples, resulting in a frequency resolution of 0.5 Hz. The Fisher z transform was used to normalize the coherence data and perform parametric statistical tests. In cats, rodents, and humans, the OB has shorter projections to the Pfr (that corresponds to orbitofrontal cortex in primates) than the Pfdl, while the Pfdl and Pp cortices have significant direct bidirectional connections (Cansler and Wesson., 2021; Zhou et al., 2019; Castro-Zaballa, 2018; Scannell et al., 1995). Hence, we focused the connectivity analysis between OB and Pfr (in 3 animals), as well as between Pfdl and Pp cortices (in 4 animals).

The same time windows were used to process the power spectrum, also using the COHER 1S script in Spike 2. The power or z-coherence values in the 100-second epochs for each EEG channel or derivative were then averaged across the different behavioral states and drug treatments.

This study also involved the use of finite impulse response filters (with a bandpass of 102-120 Hz) to obtain the HFO envelopes. This was done by filtering the data and then calculating the root mean square. A phase correlation analysis was also performed between the HFO envelopes of the OB and Pfr, including both amplitude correlations and weighted phase lag index (wPLI). The wPLI is a statistical method used to identify non-zero phase lag interdependencies between EEG time series data from pairs of electrodes, estimating the phase relationship for a specific frequency (Rawal, 2011). It measures the degree to which the phase leads and lags between the signals are non-equiprobable, regardless of the magnitude of the phase differences. In other words, it provides information about the phase consistency of the signals over time (Vinck et al., 2011). The wPLI is an extension of the phase lag index, in that takes into account the magnitude of the imaginary component of the cross-spectrum in addition to the phase relationships between signals (Vinck et al., 2011). Correlation analysis of the HFO envelopes and the wPLI of the raw recordings were performed in the derivatives constructed between the OB and Pfr (based on data from 3 animals). The 100-second analysis period was divided into 10 windows of 10 seconds, resulting in a total of 1440 windows (120 windows × 12 recordings).

The study also used wavelet transforms to better identify and visualize HFO in both time and frequency domains. The Morlet wavelet was chosen since it is well-suited for EEG analysis (Roach and Mathalon, 2008). The wavelet transforms were performed using the ND Tools Toolbox and custom MATLAB routines.

#### 2.4.2. HFO and respiratory signal

Cross-correlation frequency (CCF) maps were created between the respiration signal and the envelopes of EEG at different frequencies. The CCF-maps were obtained from 300-second windows for OB, Pfr, Pfdl, and Pp cortex in the three animals with OB and Pfr electrodes. The raw recordings were processed to generate band-pass filtered signals using the MATLAB eegfilt function (Delorme and Makeig, 2004). A 10-Hz bandwidth and 5-Hz step size were used, covering a range from 15 Hz to 150 Hz. A raster plot of CCFs was then calculated between respiration and the envelopes of each filtered signal using the built-in MATLAB xcorr and Hilbert functions.

The relationships between the phase of the respiratory wave and the amplitude of high-frequency band envelopes (phase-amplitude coupling) were analyzed using the modulation index (MI) designed by (Tort et al., 2010). The MI measures the deviation of the empirical phase-amplitude distribution from a uniform distribution, quantifying the phase-amplitude coupling.

Every MI value from each temporal window was compared to the MI values of 250 randomly generated recordings, obtained by dividing the window into small blocks and randomizing them. A permutation test was performed between the MI from the recordings and the group of randomized recordings (surrogates), and a significance level of p < 0.0001 was considered. This confirms that the MI values obtained were not the result of chance.

#### 2.4.3. Granger causality

The average direction of information flow was evaluated in the three animals with OB and Pfr channels using Granger causality (GC). GC is a statistical hypothesis test that determines whether one time series can forecast or predict the other. It is used to measure the strength and direction of the temporal relationship between two time series (Hunt et al., 2019).

GC was calculated in 500-second periods and 10-second time blocks between the OB and Pfr derivatives. The predominant direction of information flow was then obtained by subtracting the GC values in each direction: OB to Pfr (feedforward), or Pfr to OB (feedback). These calculations were performed using the MVGC multivariate GC toolbox and self-built MATLAB routines (Barnett and Seth, 2014). The significance of each 500-second period was tested using the theoretical null distribution and adjusting for multiple hypotheses.

#### 2.4.4 Statistical analyses

Data are presented as mean ± standard error (SE). The differences in HFO power, coherence, wPLI, and HFO envelope correlations, MI values (between the HFO and respiration) among states were analyzed through one-way ANOVA and Tamhane *post hoc* tests.

We compared the main direction of the information flow (calculated by the GC) between wakefulness and after ketamine using a two-tailed unpaired Student’s t-test. With the same test, we also compared the effect of nostrils occlusions on HFO power 60 seconds before and after the occlusions. Finally, the effects of the AS on HFO power were compared using two-tailed unpaired Student’s t-test before (300-second windows) and throughout the stimulus (300-second windows) during wakefulness and after ketamine administration. For all the comparisons, the null hypotheses were rejected with a significance level of p < 0.05.

## 3. Results

### 3.1. Ketamine induces HFO in the olfactory bulb and neocortex

The placement of cortical electrodes is shown in Figure 1A. Figures 1B and C illustrate examples of respiratory and EEG recordings from the OB and neocortex during wakefulness and after ketamine administration. During wakefulness, bursts of low-gamma oscillations (30 - 45 Hz) dominated the EEG signal (arrows). Ketamine injection produced a clear effect in the EEG that was associated with the emergence of HFO bursts of 200-500 ms, which occurred approximately 1 min following the injection (see “Transition to ketamine” in Figure 1B, C, arrowheads). In Figure 1D, we also show the wavelet spectrograms of the OB recording, where low-gamma and HFO were most pronounced (wavelet spectrograms from neocortical recordings are shown in Supplementary Figure 1). Notably, in the transition from wakefulness to ketamine, bursts of low-gamma and HFO occurred at different phases of the respiratory rhythm, expiration and inspiration, respectively. During the full effect of ketamine (approximately 5 minutes after the injection), the pattern of low-gamma activity became disorganized (bottom panel in 1B and D; see also Castro-Zaballa et al. 2019).

### 3.2. Ketamine increases HFO power

We compared the power of HFO in the OB and neocortical areas during wakefulness, NREM sleep, REM sleep and after ketamine injection (Figure 2A). In all these regions, ketamine induced a distinctive and prominent peak in the HFO power (100-120 Hz), indicative of a genuine neural oscillation (Yuval-Greenberg and Heeger, 2013). HFO power in the OB was higher after ketamine compared to the physiological states (Supplementary Figure 2 illustrates the whole spectrum across states). The power values were: ketamine, 95.2 ± 15.7 μV^2^; wakefulness, 12.7 ± 1.3 μV^2^; NREM, 4.7 ± 0.7 μV^2^; REM, 7.7 ± 0.9 μV^2^. The results of the statistical analysis showed that the difference between ketamine and the other states was significant (F _(4,140)_ = 380, p < 0.00001; *post hoc* ketamine Vs. all, p < 0.00001; also, wakefulness Vs. NREM, p < 0.05).

**Figure 2.**
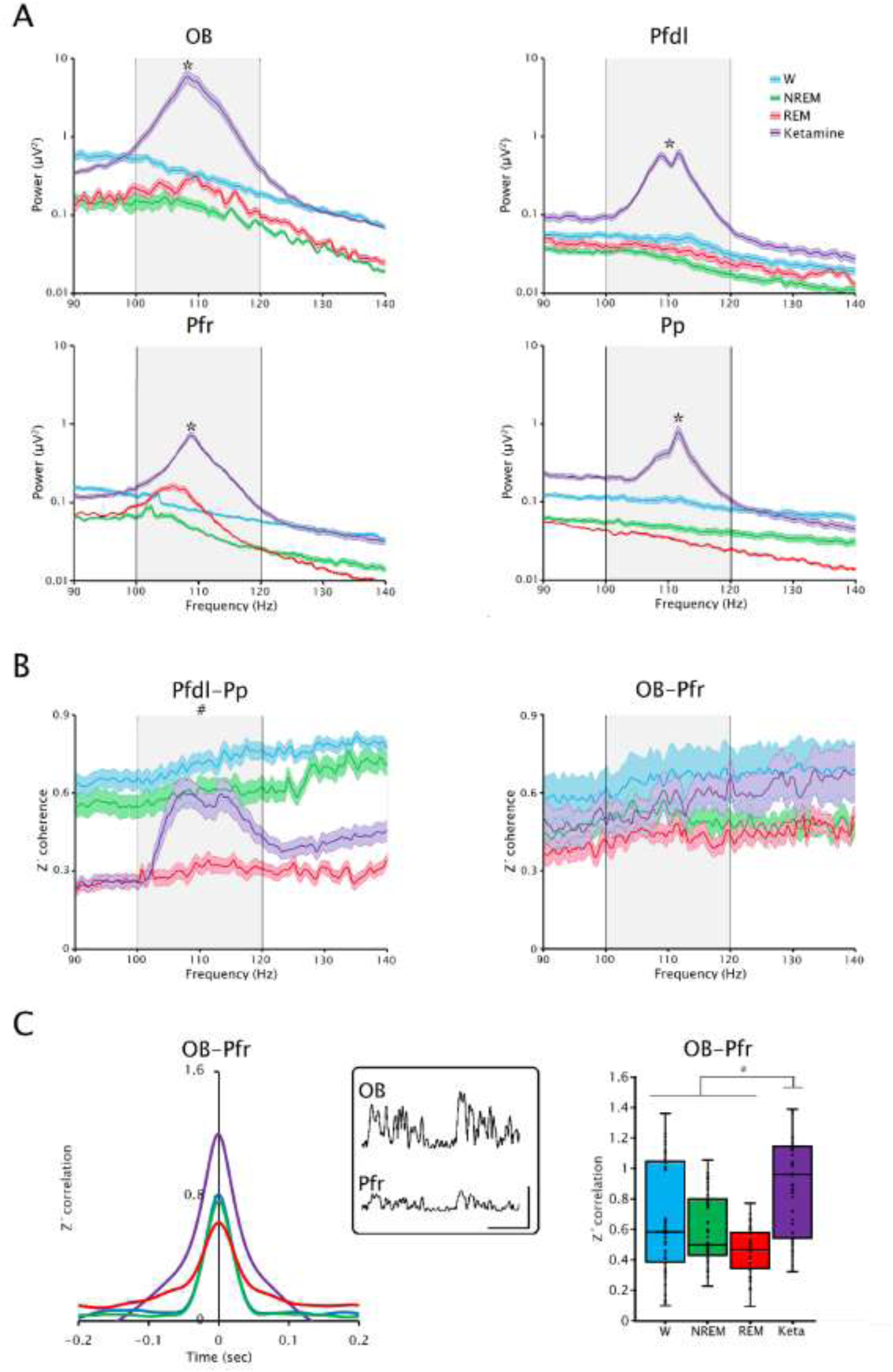
Spontaneous and ketamine-enhanced HFO power and connectivity. A. Power spectra mean ± SE from OB, Pfr, Pfdl and Pp. The gray shade denotes the HFO band. * p < 0.00001, ketamine Vs. all physiological states. B. źcoherence mean ± SE of the Pfdl-Pp and OB-Pfr derivatives; # p < 0.00001, ketamine Vs. REM sleep. C. Cross-correlation function between HFO envelopes of OB and Pfr areas (left), and box plot of linear correlations values of the HFO envelopes from the same derivative (right); * p < 0.00001, ketamine Vs. all physiological states. The inset shows the HFO envelopes of simultaneous recordings from OB and Pfr after ketamine. Inset calibration bars: 1 sec, 25 µV.

Ketamine also led to a significant increase in Pfr HFO power. The power values were: ketamine, 11.8 ± 0.6 μV^2^; wakefulness, 3.2 ± 0.1 μV^2^; NREM, 1.8 ± 0.1 μV^2^; REM, 3.3 ± 0.2 μV^2^. The statistical analysis showed that the difference between ketamine and the rest of the states was significant (F _(4,140)_ = 540, p < 0.00001; *post hoc* ketamine Vs. all, p < 0.00001; also, wakefulness Vs. NREM, p < 0.05). During REM sleep, a smaller HFO peak is visible; the power of the HFO band was significantly higher than in NREM sleep (p < 0.05) but not comparing to wakefulness.

Figure 2A also illustrates that the HFO power in the Pfdl was significantly higher under ketamine compared to the other states; the power values were: ketamine, 10.7 ± 1.0 μV^2^; wakefulness, 1.9 ± 0.2 μV^2^; NREM, 1.1 ± 0.1 μV^2^; REM, 1.4 ± 0.2 μV^2^. Again, the results of the statistical analysis showed that the difference between ketamine and the other states was significant (F _(4,188)_ = 1880, p < 0.00001; *post hoc* ketamine Vs. all, p < 0.00001; also wakefulness Vs NREM and REM, p < 0.05). Of note, even though each animal has only one HFO peak, a double peak is visible in the average power spectrum because the HFO frequency slightly differed among cats. Finally, ketamine also produced similar results in the Pp cortex; the power values were: ketamine, 11.5 ± 1.3 μV^2^; wakefulness, 3.8 ± 0.3 μV^2^; NREM, 1.8 ± 0.2 μV^2^; REM, 1.2 ± 0.03 μV^2^ (F _(4,188)_ = 1580, p < 0.00001; *post hoc* ketamine Vs. all, p < 0.00001; wakefulness Vs. NREM and REM, p < 0.05; NREM Vs. REM, p < 0.05).

### 3.4. Ketamine increases HFO coupling between cortical areas

We next examined HFO coupling between cortical areas. HFO z-coherence between Pfdl and Pp showed a synchronization peak after ketamine but not during wakefulness or sleep (Figure 2B). HFO z-coherence values were: ketamine, 0.5 ± 0.05; wakefulness, 0.72 ± 0.03; NREM, 0.60 ± 0.04; REM, 0.30 ± 0.04. Statistical analysis showed significant differences among groups (F _(4,188)_ = 154, p < 0.00001). However, HFO z-coherence after ketamine was only significantly higher than during REM sleep (*post hoc*, ketamine Vs. REM sleep, p < 0.00001; also, wakefulness Vs. NREM and REM sleep, p < 0.05 and 0.00001, respectively; NREM Vs. REM, p < 0.00001).

Although there was a large HFO power in the OB and Pfr under ketamine, the z-coherence between these areas was not different from wakefulness or sleep (ketamine, 0.56 ± 0.06; wakefulness, 0.65 ± 0.09; NREM, 0.51 ± 0.05; REM, 0.43 ± 0.04; F _(4,140)_ = 0,78, p = 0.53) (Figure 2B, right). To better understand this result, we performed a wPLI analysis. Compared to physiological states, there was no increase in HFO phase-locking under ketamine between the OB and Pfr (Supplementary Figure 3). Despite this, ketamine was associated with an increase in the HFO envelope correlation between the OB and Pfr, as can be observed in the cross-correlation function (Figure 2C, left) and in the linear correlation analysis (Figure 2C, right). Linear correlation was larger under ketamine compared to physiological states (ketamine, 0.90 ± 0.04; wakefulness, 0.67 ± 0.05; NREM, 0.60 ± 003; REM, 0.47 ± 0.02; F _(4, 1440)_ = 21.5, p < 0.00001; *post hoc* ketamine Vs. all, p < 0.00001). The strong amplitude coupling of the HFO envelopes can be readily observed in the inset of Figure 2C.

### 3.5. HFO are coupled to respiration

In our first set of results shown in Figure 1, there is a clear association between HFO and respiration. Next, we further analyzed this relationship using CCF maps and the MI (Figure 3), which measures the correlation between the phase of respiration and the amplitude of HFO.

**Figure 3.**
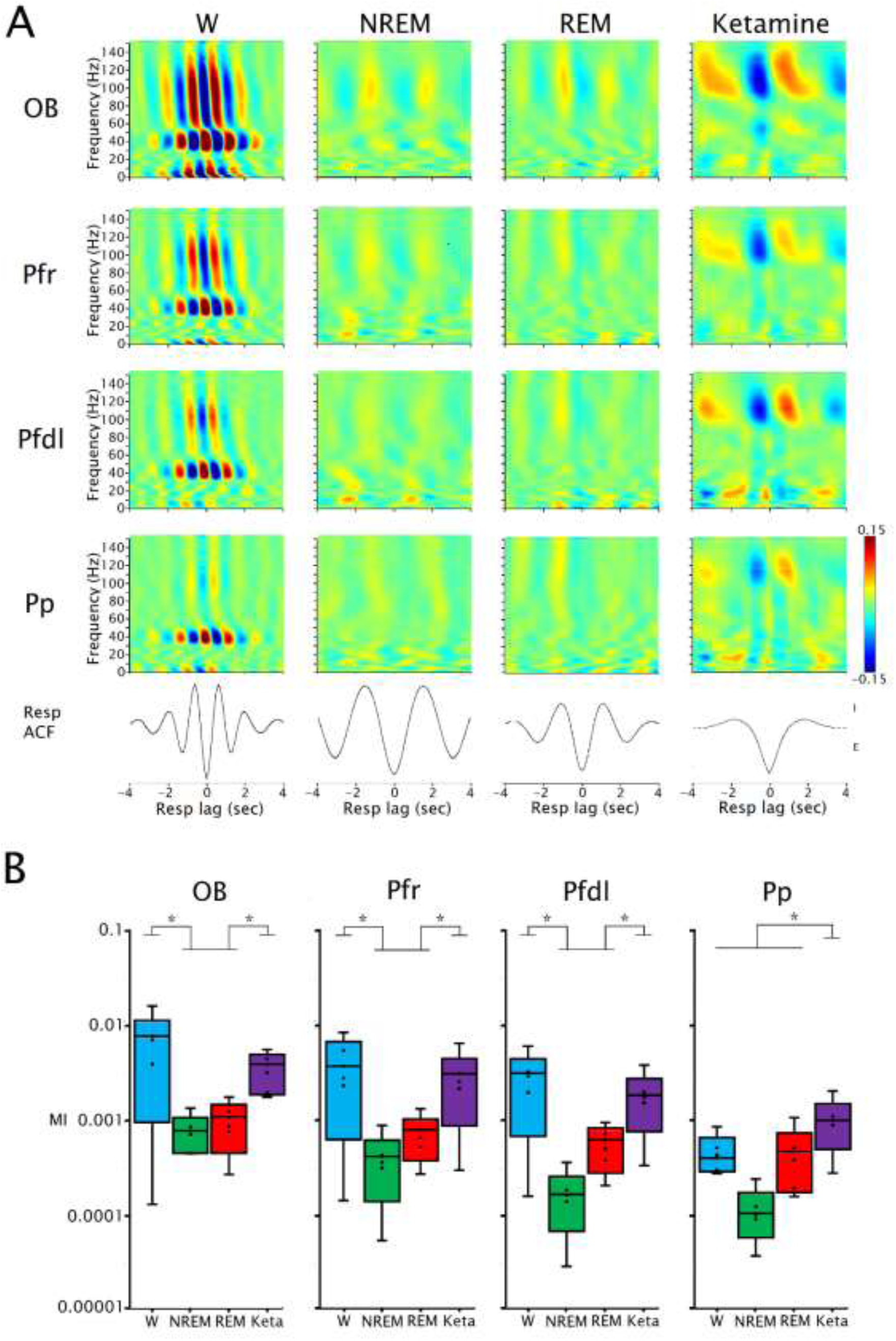
Spontaneous and ketamine-enhanced HFO are in phase with respiration. A. Cross-correlation frequency maps between the respiratory signal and the amplitude envelope of the EEG signals from the olfactory bulb (OB), rostral prefrontal (Pfr), dorsolateral prefrontal (Pfdl) and parietal posterior (Pp) cortices of a representative cat. The signals were filtered between 2 and 150 Hz (10-Hz bandwidth and 5-Hz steps) during wakefulness (W), NREM sleep, REM sleep and after ketamine. Bottom, autocorrelation function (ACF) of the respiratory signal. E, expiration, I, inspiration. B. Box plot of the modulation index (MI) values for OB, Pfr, Pfdl and Pp during W, NREM, REM and under ketamine (Keta). The ANOVA analysis of the MI for the OB was F _(4,140)_ = 28; Pfr, F _(4,140)_ = 25; Pfdl, F _(4,140)_ = 24 and Pp, F _(4,140)_ = 18; *, p < 0.05.

Figure 3A shows in a representative cat that HFO amplitude correlates with the respiratory cycle both during wakefulness and under ketamine. This was marked in the OB but still present in neocortical areas. The CCF map of the transition from wakefulness to ketamine is shown in Supplementary Figure 4.

Group-level analysis of phase-amplitude coupling confirmed the association between nasal respiration and HFO, and showed that the MI values were significantly above chance in these areas both during wakefulness and ketamine (compared with surrogates, data not shown). HFO coupling to respiration was below chance levels during NREM and REM sleep (even in OB, which showed weak HFO power peaks during NREM and REM sleep). Finally, we compared the magnitude of MI values among states and found that HFO coupling in the OB, Pfr and Pfdl was larger during wakefulness and ketamine compared to sleep, while in the Pp MI values were larger after ketamine compared to the other states (Figure 3B).

Interestingly, ketamine-enhanced HFO were coupled to the respiratory cycle on a longer time scale. This is due to the decrease in respiratory frequency caused by the drug (see the respiratory autocorrelation functions - ACF-in Figure 3A). The respiratory frequency was different among states, with the following mean values: wakefulness, 0.44 ± 0.02 Hz in; NREM sleep, 0.32 ± 0.01; REM sleep, 0.40 ± 0.04; ketamine, 0.23 ± 0.01. Statistical analysis showed that the respiratory frequency under ketamine was significantly lower than all other states (F _(4,188)_ = 18.5, p < 0.00001; *post hoc* ketamine Vs. all, p < 0.00001).

### 3.6. The information flow of ketamine-enhanced HFO is from OB to neocortex

The coupling of HFO with the inspiratory phase of respiration suggests that HFO originates within the OB and spreads towards the neocortex. To test this hypothesis, we performed GC analysis to estimate the direction of information flow between the OB and Pfr.

Figure 4 shows the GC spectra for the OB→Pfr (feedforward, FF) and Pfr→OB (feedback, FB) directions, as well as the difference between them. These results showed a strong causality for the HFO band in the OB→Pfr (FF) direction under ketamine, but not during wakefulness. The result of subtracting FB minus FF after ketamine (−0.27 ± 0.07) was different compared to wakefulness (0.02 ± 0.02; T = 16, p < 0.00001). On the other hand, GC analysis did not show a clear HFO directionality between neocortical areas neither during wakefulness nor after ketamine (Supplementary Figure 5).

**Figure 4.**
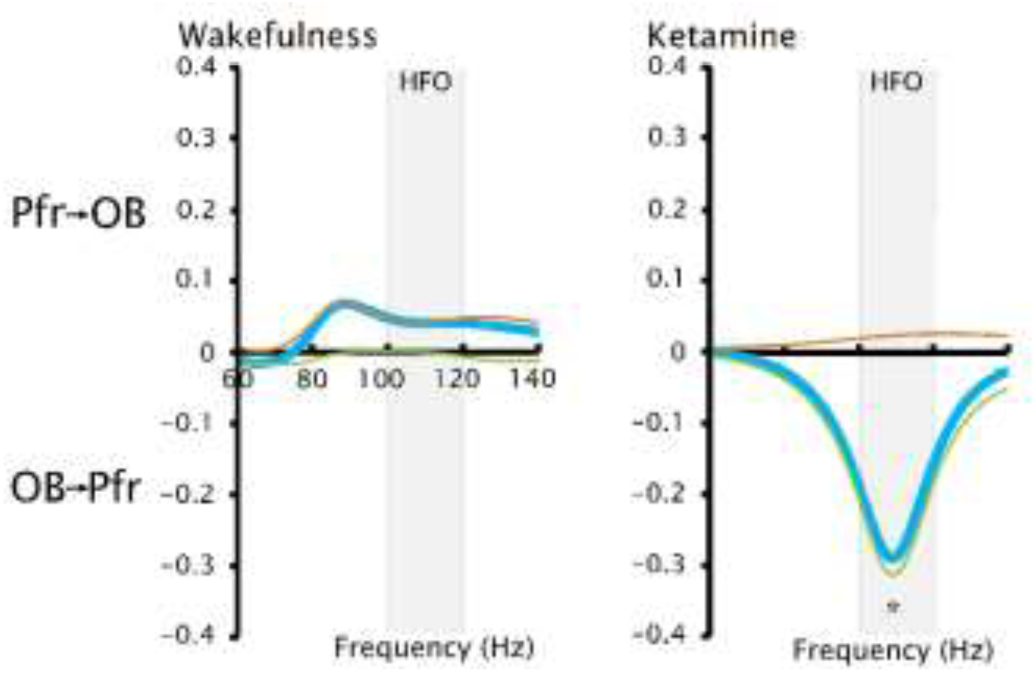
The OB drives ketamine-enhanced but not spontaneous HFO. Mean Granger causality spectra of HFO for in the Pf→OB (FB, red) and OB→Pf (FF, green) directions and their subtraction (sky blue), during wakefulness and under ketamine. The gray shade denotes the HFO band. The result of subtracting FB minus FF after ketamine was different compared to wakefulness, * p < 0.00001.

### 3.7. Ketamine-enhanced HFO are dependent on nasal airflow

To test if HFO depend on the airflow through the nostrils, we performed naris occlusion experiments during wakefulness and under ketamine. Occluding the ipsilateral (right) or both nostrils increased HFO power in the OB during wakefulness (Figure 5A). However, after ketamine, occluding the ipsilateral or both nostrils greatly reduced HFO power. In the Pfr, while nostril occlusions resulted in small increases in HFO during wakefulness, both ipsilateral and bilateral occlusions produced significant decreases in HFO after ketamine. Quantitative bar graphs and their statistics are shown in Figure 5B. These experiments demonstrate that ketamine-enhanced HFO depend on nasal airflow, are likely to originate in the OB, and are integrated bilaterally in the Pfr cortex.

**Figure 5.**
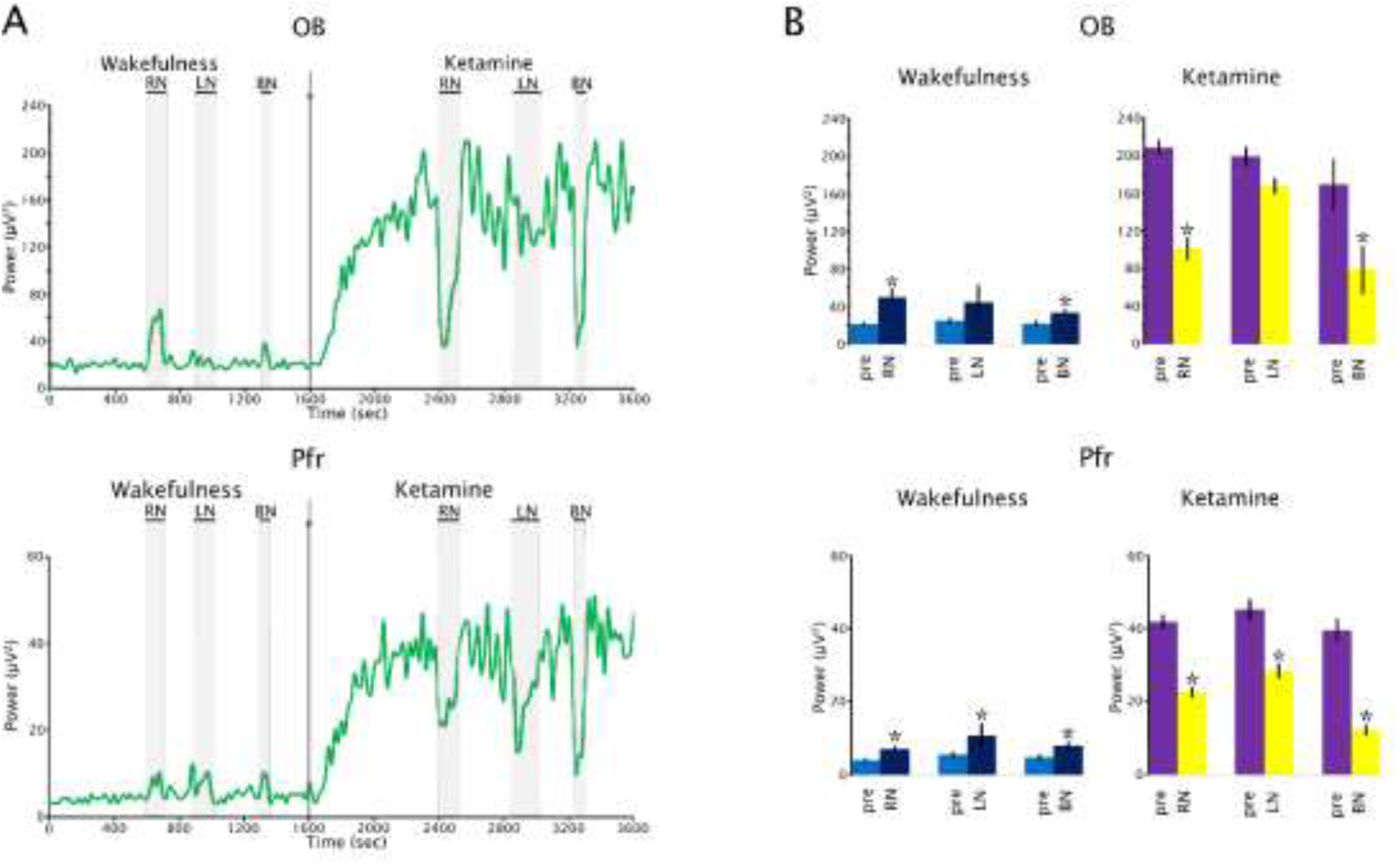
Nasal occlusion reduces the power of ketamine-enhanced HFO. A. Effect of nasal occlusion on HFO power in the olfactory bulb (OB) and prefrontal (Pfr) cortex before (during wakefulness) and following the injection ketamine in a representative animal. The gray shade areas represent nasal occlusion: RN, right nasal occlusion; LN, left nasal occlusion; and BN, bilateral nasal occlusion. B. Mean ± SE of the HFO power of OB and Pfr during wakefulness and after ketamine, before the nasal occlusion (pre) and under the effect of nasal occlusion; * p < 0.01.

### 3.8. HFO are not modified by auditory stimulation

Figure 6 shows representative spectrograms and the amplitude of the HFO-envelopes during baseline and ketamine recordings. Wakefulness was associated with high low-gamma power which augments as the level of alertness increases induced by AS, while HFO power remains low (Figure 6A, top). This also can be observed in the amplitude of the envelopes of low-gamma and HFO oscillations (Figure 6A, bottom) (see also Supplementary Table 1 for statistical analysis). Upon administration of ketamine, there is a marked increase in HFO power (Figure 6B, top) and amplitude of the HFO envelopes (Figure 6B. bottom). However, after ketamine AS affect neither low-gamma nor HFO (Figure 6 B; Supplementary Table 1).

**Figure 6.**
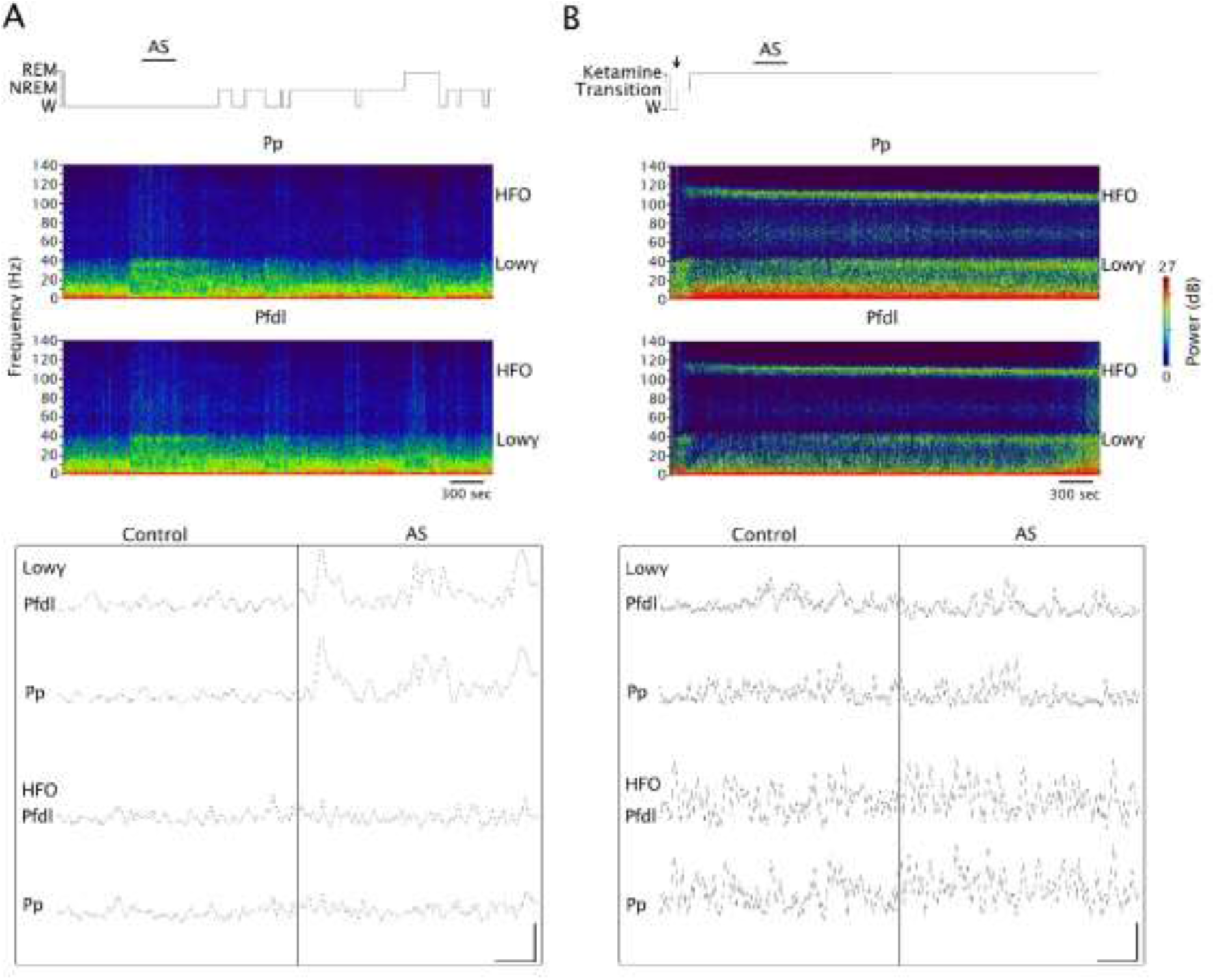
Auditory stimulation does not affect HFO. A. Top. Hypnogram and power spectrograms of prefrontal (Pfdl) and posterior-parietal (Pp) cortices during a baseline recording of a representative animal. AS indicates the auditory stimulation. Bottom. Amplitude envelopes of low-gamma and HFO during wakefulness before AS (control) and during AS. B. The same analyses are shown after ketamine administration. The arrow indicates the moment when ketamine was injected. Transition, transition to ketamine; W, wakefulness. Calibration bars: 0.5 sec, 10 µV.

## 4. Discussion

In the present report, we studied cortical HFO in cats during wakefulness, sleep, and after a subanesthetic dose of ketamine. During wakefulness, although we did not identify a discernable HFO peak in the power spectrum, we found a relationship between HFO bursts and respiration. Ketamine significantly increased HFO power in both the OB and neocortex. Further analysis revealed that this rhythm was coupled to respiration, propagated from the OB to the neocortex, and was reduced by nasal airflow occlusion.

### 4.1. HFO during wakefulness and sleep

Previous works in rats have identified HFO during wakefulness and REM sleep in several brain regions, including neocortex, hippocampus, amygdala, striatum, subthalamic nucleus, and substantia nigra (Buzsáki et al., 1983; Cordon et al., 2015; Kealy et al., 2017; Nicolás et al., 2011b; Ye et al., 2018). However, our results in cats showed that HFO during wakefulness and sleep, although present, are not visible as a discernable peak in the power spectra.

We also found that HFO coherence was the weakest during REM sleep. Thus, both HFO and low-gamma (≍40 Hz) coherence are very low during REM sleep in the cat (Castro et al., 2014, 2013; Castro-Zaballa et al., 2019b; Cavelli et al., 2017; Torterolo et al., 2016). This fact differs from previous studies in rats, where high intra-hemispheric and low inter-hemispheric synchronization of the HFO was observed (Cavelli et al., 2018).

### 4.2. HFO are enhanced by subanesthetic administration of ketamine

In accordance with our findings in cats that ketamine increases HFO power in the OB and neocortical areas, rodent studies have reported that NMDA receptor antagonists increase HFO power in several brain regions, such as neocortex, hippocampus, thalamus, substantia nigra, caudate, accumbens and subthalamic nucleus (Cordon et al., 2015; Flores et al., 2015; Hunt et al., 2011; Kulikova et al., 2012; Nicolás et al., 2011b; Phillips et al., 2012; Caixeta et al., 2013; Olszewski et al., 2013; Hunt and Kasicki, 2013; Hunt et al., 2006). This likely reflects an increment in local synchronization of fast neuronal activity (Cavelli et al., 2017).

In rats, NMDA receptor antagonists also increase HFO coupling in the cortico-basal ganglia circuit (Cordon et al., 2015), as well as between OB and the ventral striatum (Hunt et al., 2019). In the present study, we found that after ketamine there is a peak in HFO neocortical interregional coherence, with values that were higher than during REM sleep, indicating larger long-distance neocortical coupling after ketamine compared to REM sleep (Cavelli et al., 2017). In contrast, our results suggest that ketamine increased the coupling between the OB and Pfr in terms of amplitude, but not in phase. Further, this increase was reflected as an increment in the correlation between OB and Pfr envelopes of HFO (Figure 2C).

### 4.3. HFO are modulated by respiration and propagated from the OB

As shown previously (Cavelli et al., 2020; González et al., 2023a, 2023b), during wakefulness, a significant correlation was observed between respiration and low-gamma activity in the OB, piriform cortex and neocortical regions, which declined during sleep. Interestingly, a strong correlation was also observed in HFO during wakefulness but not during sleep, highlighting the state-dependency of these oscillations. Also a gradient of correlation between HFO and respiration was observed across the brain during wakefulness, with the OB having the strongest and Pp the weakest correlation, indicating a greater impact of respiration on anterior brain regions (Tort et al., 2018). Notably, a similar pattern was seen under ketamine, with a stronger coupling in the OB and minimal coupling in the Pp. However, the coupling with the respiration under ketamine was restricted to HFO, while during wakefulness also included lower frequencies (Figure 3).

HFO and low-gamma bands coupled differentially with the respiratory cycle. The low-gamma band was linked to the expiratory phase (Cavelli et al., 2020), while HFO were linked to the inspiratory phase both during wakefulness and after ketamine. This relationship was also observed in rodents (Wróbel et al., 2020), where HFO after ketamine in the OB, ventral striatum, and prefrontal cortex were found to be coupled to inspiration.

To test the hypothesis that nasal respiration is crucial for HFO in cats, we conducted nasal occlusion experiments. Nasal occlusion after ketamine reduced the power of HFO, a finding consistent with (Wróbel et al., 2020) who found that unilateral nasal occlusion in rats reduced HFO power only in the ipsilateral side in the OB, ventral striatum, and prefrontal cortex. Similarly, in accordance with our findings, Średniawa et al., (2021) found that HFO in the OB of cats under ketamine-xylazine anesthesia is driven by nasal airflow and is coupled with nasal respiration; unilateral naris blockade also reduced HFO power on the same side of the OB, which quickly recovered once the blockade was removed.

Together, these results demonstrate that nasal airflow drives ketamine-enhanced HFO in multiple brain regions, with the OB playing a crucial role in the dissemination of this rhythm. This is supported by our GC analyses, which measures the direction of information flow. We found that under ketamine, HFO flow of information is from the OB towards the Pfr cortex, in a feedforward direction. This is also in line GC analyses in rodents showing ketamine-dependent HFO flows from the OB to the ventral striatum (Hunt et al., 2019). This directionality was not observed during wakefulness.

### 4.4. Differences between HFO and the gamma band

In cats, low-gamma and HFO recorded in the EEG differ in important ways: 1. Low-gamma (≍ 40 Hz) power and coherence vary across different behavioral states (Castro et al., 2014, 2013; Cavelli et al., 2017, 2015; González et al., 2019), while the state-dependency of HFO is subtler, only revealed when the coupling with the respiration is studied. 2. Ketamine increases low-gamma power in the neocortex but reduces gamma coherence (Castro-Zaballa et al., 2019b). In contrast, injection of ketamine is associated with a prominent 100-120 Hz power in the spectrum; a narrow-band peak in coherence can also be observed between neocortical derivates. 3. During wakefulness, low-gamma activity is synchronized with the expiration phase of respiration, while HFO are associated with the inspiration phase. Under ketamine, the synchronization of low-gamma activity with respiration is absent. 4. AS during wakefulness increases alertness and results in a significant increase in low-gamma power and coherence (Castro et al., 2014, 2013; Castro-Zaballa et al., 2019b, 2019a; Torterolo et al., 2016). However, AS does not affect HFO during wakefulness nor after ketamine, suggesting that HFO are not primarily associated with processing novel auditory information. These findings emphasize that low-gamma activity and HFO represent conceptually different phenomena. A recent study by (Cichon et al., 2023) in mice found that in several cortical layers and regions, ketamine causes spontaneously active neurons to become suppressed, while previously silent neurons become activated. It would be interesting to explore if this phenomenon has a link with the switch of low-gamma predominance during wakefulness to HFO predominance after ketamine.

### 4.5. HFO in cognition and relationship with psychosis

Clinical and preclinical findings have implicated NMDA receptors in the pathophysiology of psychotic disorders (Kantrowitz and Javitt, 2012; Lahti et al., 1995; Schwartz et al., 2012). Indeed, *post-mortem* studies of patients with schizophrenia have reported a decrease in NMDA receptors in the dorsolateral prefrontal cortex and hippocampus (Konradi and Heckers, 2003).

HFO are prominent in pharmacological models of psychosis (Caixeta et al., 2013; Cordon et al., 2015; Hunt et al., 2019; Hunt and Kasicki, 2013; Olszewski et al., 2013; Średniawa et al., 2021; Wróbel et al., 2020). Although most studies investigating ketamine-dependent HFO have used rodents, here we advanced these studies using cats, which have a gyrencephalic anatomy similar to humans. Indeed, a recent scalp EEG study in humans has shown increases in broadband HFO in midline areas after subanesthetic ketamine and d-cycloserine, suggesting that HFO data from experimental animals could be translatable to humans (Nottage et al., 2023). In addition, available evidence suggests that schizophrenia is associated with abnormal 60-120 Hz oscillations which could potentially explain some core features of the disorder, such as cognitive impairments (Uhlhaas and Singer, 2013).

REM sleep may be considered a natural model of psychosis (Hobson, 1997, 2009). While long-distance gamma coupling in neocortex is reduced in both REM sleep (Castro et al., 2013) and ketamine models of psychosis, HFO coupling is reduced in REM sleep but not under ketamine. Hence, the profile of HFO differs between both model of psychosis.

### 4.6. Technical considerations

To date, spontaneous and ketamine-induced HFO were almost exclusively studied in rodents, with few studies in more translatable models. Here, the recordings in cats were obtained in a semi-restricted condition, in which postures or movements did not influence the recordings, thereby reducing the possibility of artifacts.

We used 15 mg/kg of ketamine, which when administered as a single agent is subanesthetic; the anesthetic dose in cats is ≥ 25 mg/kg (Arnbjerg, 1979; Hanna et al., 1988). The typical behavioral syndrome (hyperlocomotion, ataxia signs and stereotypes) that develops in rodents following subanesthetic doses of ketamine was not observed in cats. On the contrary, five minutes following the injection in freely-moving cats, the animals remained lying on the floor unable to stand up, but responded to sound stimulus by directing their gaze toward the sound source. In the absence of stimuli, the cats moved their head from one side to another (probably responding to hallucinatory activity). The animals retained muscular tone, showed hyper-salivation, and dilated pupils. Usually, around 40 minutes after ketamine administration the animals tried to stand up, and three hours later were fully recovered (Castro-Zaballa et al., 2019b).

## 5. Conclusions

In the present report, we show in the cat that cortical HFO are present during wakefulness, are reduced in power during both NREM and REM sleep, and enhanced by a subanesthetic dose of ketamine. This rhythm was larger in the OB and frontal cortical regions, and dependent on nasal respiration. Aberrant HFO induced by ketamine across cortical areas would be expected to disrupt information processing and may account for some of the psychoactive effects produced by ketamine.

## Acknowledgements

This research was supported by the following grants: CSIC-I+D grupos 2022-group ID-22620220100148, and CSIC-I+D-2020-393.

## CRediT (Contributor Roles Taxonomy) author statement

**Santiago Castro-Zaballa:** Conceptualization, Methodology, Validation, Formal analysis, Investigation, Data Curation, Writing-Original draft preparation, Visualization, Project administration, Funding acquisition.

**Joaquín González**: Software, Validation, Writing - Review & Editing, Visualization.

**Matías Cavelli**: Methodology, Software, Validation, Writing - Review & Editing.

**Diego Mateos**: Validation, Formal analysis, Writing - Review & Editing.

**Claudia Pascovich**: Validation, Writing - Review & Editing.

**Adriano Tort**: Validation, Writing - Review & Editing.

**Mark Jeremy Hunt**: Methodology, Conceptualization, Validation, Writing - Review & Editing.

**Pablo Torterolo**: Conceptualization, Methodology, Validation, Writing - Review & Editing, Visualization, Supervision, Project administration, Funding acquisition.

Declarations of interest: none

## Notes

### Competing Interest Statement

The authors have declared no competing interest.

